# Remodelling of carbon metabolism during sulfoglycolysis in *Escherichia coli*

**DOI:** 10.1101/2022.09.09.507388

**Authors:** Janice W.-Y. Mui, David P. De Souza, Eleanor C. Saunders, Malcolm J. McConville, Spencer J. Williams

## Abstract

Sulfoquinovose (SQ) is a major metabolite in the global sulfur cycle produced by nearly all photosynthetic organisms. One of the major pathways involved in the catabolism of SQ in bacteria, such as *Escherichia coli*, is a variant of the glycolytic Embden-Meyerhof-Parnas (EMP) pathway termed the sulfoglycolytic EMP (sulfo-EMP) pathway, which leads to consumption of three of the six carbons of SQ and excretion of 2,3-dihydroxypropanesulfonate (DHPS). Comparative metabolite profiling of aerobically Glc-grown and SQ-grown *E. coli* was undertaken to identify the metabolic consequences of switching from glycolysis to sulfoglycolysis. Sulfoglycolysis was associated with the diversion of triose-phosphates to synthesize sugar phosphates (gluconeogenesis), and an unexpected accumulation of trehalose and glycogen storage carbohydrates. Sulfoglycolysis was also associated with global changes in central carbon metabolism, as indicated by changes in levels of intermediates in the tricarboxylic acid (TCA) cycle, the pentose phosphate pathway (PPP), polyamine metabolism, pyrimidine metabolism and many amino acid metabolic pathways. Upon entry into stationary phase and depletion of SQ, *E. coli* utilize their glycogen, indicating a reversal of metabolic fluxes to allow glycolytic metabolism.

**Importance:** The sulfosugar sulfoquinovose is estimated to be produced on a scale of 10 billion tonnes per annum, making it a major organosulfur species in the biosulfur cycle. Microbial degradation of sulfoquinovose through sulfoglycolysis allows utilization of its carbon content and contributes to biomineralization of its sulfur. However, the metabolic consequences of microbial growth on sulfoquinovose are unclear. We use metabolomics to identify the metabolic adaptations that *Escherichia coli* undergoes when grown on sulfoquinovose versus glucose. This revealed increased flux into storage carbohydrates through gluconeogenesis, and reduced flux of carbon into the TCA cycle and downstream metabolism. These changes are relieved upon return to stationary phase growth and reversion to glycolytic metabolism. This work provides s new insights into the metabolic consequences of microbial growth on an abundant sulfosugar.

## Introduction

Sulfoquinovose (SQ) is a 6-sulfonated analogue of glucose (Glc) that is produced by most photosynthetic organisms (Fig. 1A).(1, 2) SQ occurs primarily as the polar head group of sulfoquinovosyl diacylglyceride (SQDG), a major glycolipid in plant chloroplasts and cyanobacteria.(1, 3) Global production of SQ is estimated to be of the order of 10 billion tonnes per annum.(4) Consequently, SQ is a major compound in the biogeochemical sulfur cycle. While the biosynthesis of SQ and SQDG occurs within plants and cyanobacteria,(1, 5) its catabolism appears to be mediated exclusively by bacteria.

**Fig 1.**
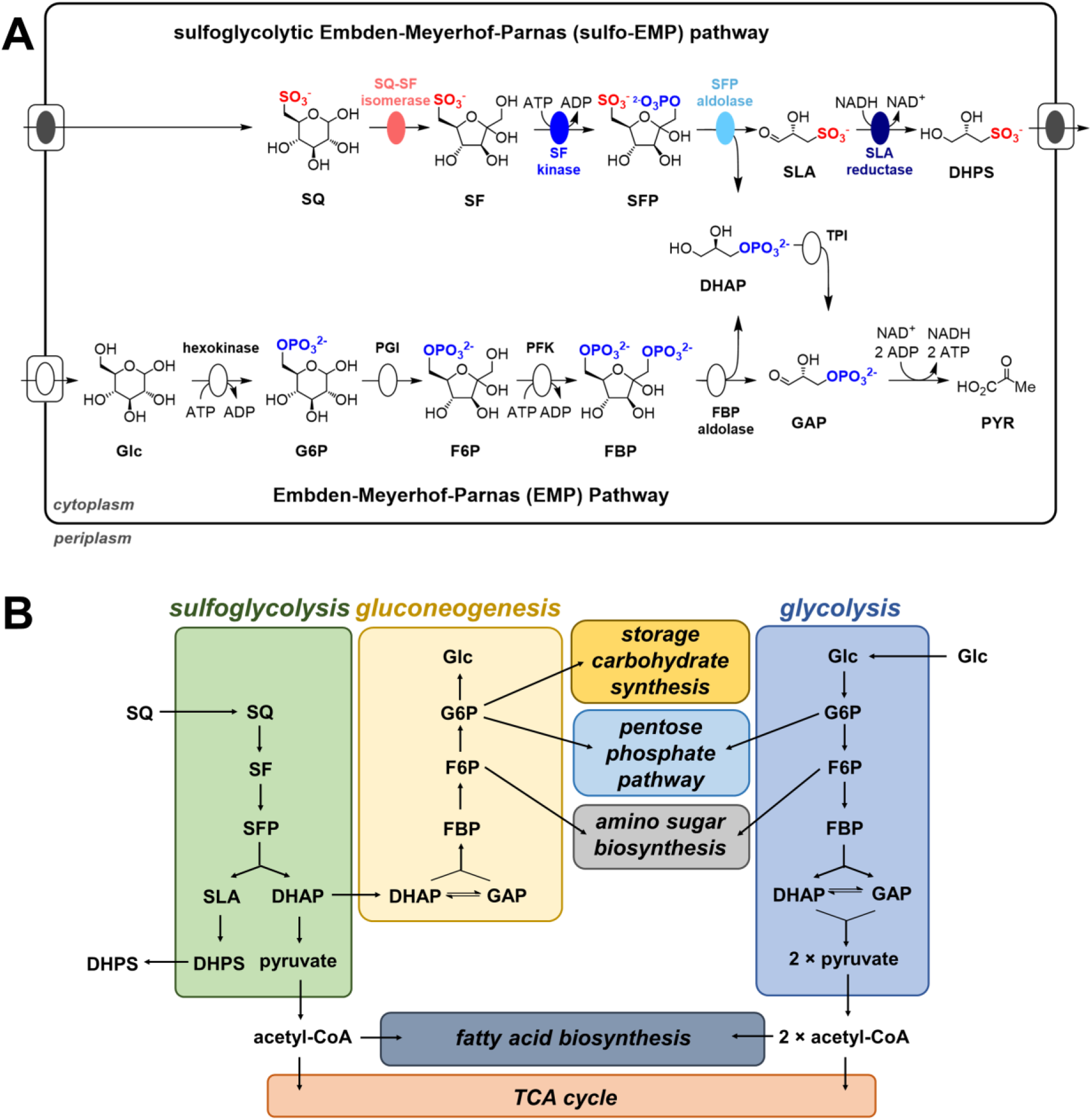
(A) The glycolytic Embden-Meyerhof-Parnas (EMP) pathway, (B) the sulfoglycolytic Embden-Meyerhof-Parnas (sulfo-EMP) pathway and (C) diagrammatic representation of central carbon metabolism in glycolytic and sulfoglycolytic *E. coli*. Glc, glucose; G6P, glucose-6-phosphate; F6P, fructose-6-phosphate; FBP, fructose-bisphosphate; GAP, glyceraldehyde-3-phosphate; DHAP, dihydroxyacetone phosphate; SQ, sulfoquinovose; SF, sulfofructose; SFP, sulfofructose-1-phosphate, SLA, sulfolactaldehyde; DHPS, 2,3-dihydroxypropanesulfonate; PGI, phosphoglucose isomerase; PFK, phosphofructose kinase 1; FBP aldolase, fructose-bisphosphate aldolase, TPI, triose-phosphate isomerase; SQ isomerase, sulfoquinovose isomerase; SF kinase, sulfofructose kinase; SFP aldolase, sulfofructose-1-phosphate aldolase; acetyl-CoA, acetyl coenzyme A; TCA cycle, tricarboxylic acid cycle.

The pathways of catabolism of SQ are termed ‘sulfoglycolysis’.(6, 7) The first sulfoglycolytic pathway was described in *Escherichia coli*,(8) and was coined the sulfoglycolytic Embden-Meyerhof-Parnas (sulfo-EMP) pathway, owing to its similarity to the ‘investment phase’ of glycolysis or the upper glycolytic EMP pathway (Fig. 1A), in which two molecules of ATP are consumed to convert Glc to two molecules of glyceraldehyde-3-phosphate (GAP). Upper glycolysis comprises the steps catalyzed by hexokinase (phosphorylation of Glc to glucose-6-phosphate (G6P)), G6P isomerase (conversion of G6P to fructose-6-phosphate (F6P)), F6P kinase (phosphorylation of F6P to form fructose-bisphosphate (FBP)), FBP aldolase (cleavage of FBP into two 3-carbon (3C) fragments: GAP and dihydroxyacetone phosphate (DHAP)), and triose-phosphate isomerase (interconversion of GAP and DHAP). During the ‘payoff phase’, or lower glycolysis, two molecules of GAP are converted to two molecules of pyruvate over five steps, producing energy as ATP and reducing power as NADH. In contrast, the sulfo-EMP pathway converts SQ to DHAP and sulfolactoaldehyde (SLA) using a dedicated SQ isomerase (converts SQ to sulfofructose (SF)), a SF kinase (phosphorylates SF to give sulfofructose-1-phosphate (SFP)), and a SFP aldolase (cleaves SFP into DHAP and SLA) (Fig. 1A). As in glycolysis, DHAP is isomerized to GAP, and further catabolized in lower glycolysis, with production of ATP and NADH, while the other 3C fragment, SLA, is reduced to 2,3-dihydroxypropanesulfonate (DHPS), generating NAD^+^, and excreted from the cell. The sulfo-EMP pathway therefore has balanced NAD^+^/NADH production, but yields half the ATP and carbon per hexose sugar compared to the glycolytic EMP pathway.(9)

*E. coli* shows a clear preference for growth on Glc versus SQ. Glc-adapted *E. coli* K-12 strain MG1655 takes two weeks to adapt to growth on SQ, and once adapted, it grows more slowly than on Glc.(8) Maximum biomass of *E. coli* grown on SQ is also approximately half of cultures grown on Glc.(8) Reduced growth and biomass on SQ could reflect the lower yield of carbon, ATP and NADH in the sulfo-EMP pathway and supply of energy and precursors for anabolic processes. The sulfo-EMP pathway also bypasses the steps in glycolysis that produce G6P or F6P, which are needed to supply precursors for the pentose phosphate pathway (PPP) and sugar nucleotide and cell wall biosynthesis, respectively (Fig. 1B). SQ-grown *E. coli* must therefore additionally divert the triose phosphates (triose-P) DHAP/GAP produced by sulfoglycolysis into G6P and F6P,(10, 11) while maintaining sufficient flux into lower glycolysis and pyruvate production to sustain the tricarboxylic acid (TCA) cycle and fatty acid biosynthesis. The switch from glycolysis to sulfoglycolysis therefore requires large-scale metabolic changes that may be responsible for the slow transition from Glc to SQ. While these changes have yet to be characterized, limited study has been done: Burrichter and coworkers provided evidence of a novel mixed-acid fermentation and conversion of SQ-derived triose-P to succinate, acetate and formate when *E. coli* was grown on SQ under anaerobic conditions.(12)

In this work, we investigate metabolic changes occurring in *E. coli* grown with Glc or SQ as sole carbon sources under aerobic conditions using comparative metabolite profiling. Unexpectedly, we find evidence for extensive diversion of SQ-derived carbon into glycogen and trehalose storage carbohydrates in SQ-grown *E. coli* during logarithmic phase, suggestive of limitation in other pathways required for growth. These storage carbohydrates are utilized following commencement of stationary phase, suggestive of a reverse diauxic shift whereby bacteria temporarily switch back to glycolysis following exhaustion of their primary carbon source. Untargeted metabolite analysis reveals further changes to metabolite levels across a range of metabolic pathways during sulfoglycolytic metabolism: these include clear perturbations to the TCA cycle, the PPP, polyamine metabolism, pyrimidine metabolism and many amino acid metabolic pathways. This work shows that metabolic adaptation to sulfoglycolytic growth in *E. coli* requires the simultaneous operation of gluconeogenesis and lower glycolysis, leading to the accumulation of storage carbohydrates. We propose that the accumulation of storage carbohydrates during SQ metabolism is associated with primary deficits in energy (as ATP), reducing power and carbon-based building blocks.

## Results

### Growth on SQ leads to major changes in *E. coli* metabolism

*E. coli* BW25113 were grown to mid-log phase on Glc (4 mM) and on SQ (8 mM) as sole carbon sources. The SQ concentration chosen was double that of Glc to account for the fact that *E. coli* excrete half of the internalized SQ (carbons 4-6 and the sulfonate group) as DHPS; this approach was also utilized by Burrichter and coworkers.(12) This doubling of SQ concentration does not appear to have substantial impact on the growth rate; growth was approximately 5-fold slower than in Glc-grown *E. coli* (compared with 3.8-fold in a previous study for *E. coli* MG1655(8)). Cells were harvested, metabolically quenched and intracellular metabolites were extracted into MeOH:H_2_O (3:1 v/v),(13) derivatised, then analysed by gas chromatography-electron ionization-triple quadrupole mass spectrometry (GC-EI-QqQ-MS).(14) Metabolites were detected by multiple reaction monitoring (521 targets, representing approximately 350 metabolites, with two MRM transitions (qualifier and quantifier) per target). Data was manually inspected and curated prior to more detailed analysis.

In total, 146 metabolites were detected in at least one growth condition, of which 117 had statistically significant differences in abundance in bacteria grown on the two carbon sources (False Detection Rate-adjusted p-value < 0.05). As shown using a volcano plot, 36 metabolites had lower abundance in SQ-grown versus Glc-grown *E. coli*, and 81 had a higher abundance (Fig. 2A). Fold changes spanned several orders of magnitude: lysine was >4000-fold less abundant, several metabolites were >100-fold less abundant and several were >10-fold more abundant in SQ-grown *E. coli* (Fig. 2B, 2C). There were also large increases in hexose-based sugars in SQ-fed *E. coli*.

**Fig 2.**
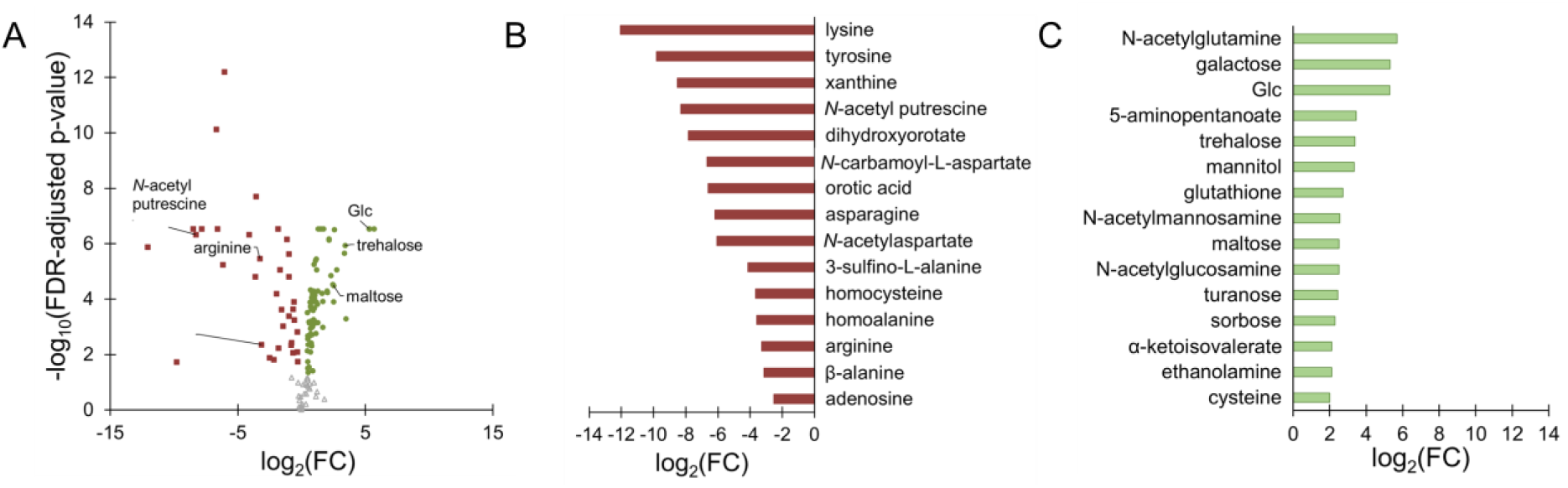
Fold change (FC) in metabolite abundance in *E. coli* grown on Glc versus SQ. (A) Volcano plot of results with selected metabolites labelled, (B) 15 largest decreases in abundance in SQ-grown *E. coli* and (C) 15 largest increases in abundance in SQ-grown *E. coli*.

### Amino acid pools are perturbed during sulfoglycolysis

Many amino acid pools underwent very large perturbations upon changing from growth on Glc to SQ (Fig. 2B, 2C, Table S1). Tyrosine was 900-fold less abundant in SQ-grown *E. coli*, yet phenylalanine and its derivatives phenyllactate and phenylpyruvate(15–17) showed no statistically significant changes (Fig. 2B, Table S1). Lysine was > 4000-fold less abundant in SQ-grown *E. coli* (Fig. 2B, Table S1). Alanine, methionine, threonine, valine, arginine and asparagine were also less abundant (Table S1). In contrast, serine, cysteine, glycine, isoleucine, leucine, glutamate, aspartate and proline were all more abundant in SQ-grown *E. coli* (Table S1).

Changes were also observed for intermediates in amino acid biosynthetic and degradation pathways. The cysteine degradation product 3-sulfino-L-alanine(15–17) was 17-fold less abundant in SQ-grown *E. coli*, while the lysine degradation products 5-aminopentanoate, glutarate, 2-hydroxyglutarate and α-ketoglutarate(15–17) and the valine degradation product 3-hydroxyisobutyrate(15–17) were all more abundant (Table S1). The leucine precursors 2-isopropylmalate and ketoleucine(15–17) and the leucine and valine common precursor α-ketoisovalerate(15–17) were more abundant in SQ-grown *E. coli*, while the valine, leucine and isoleucine common precursor α-ketobutyrate(15–17) was less abundant (Table S1).

Putrescine levels were 1.5-fold higher in mid log-phase SQ-grown versus Glc-grown *E. coli* (Fig. 3A). There was also 1.5-fold more proline and glutamate, but 10-fold less arginine in SQ-grown *E. coli* (Fig. 3A). Putrescine is derived either from arginine via agmatine (which releases urea), or from glutamate or proline via ornithine, the latter being more direct (Fig. 3C).(15–20) It has been reported that in *E. coli*, urea is almost exclusively produced by putrescine biosynthesis from arginine via agmatine.(18, 21) The observation of 3-fold less urea in SQ-grown *E. coli* may indicate lower dependence on the agmatine pathway. Putrescine has two possible fates in *E. coli*; it can be oxidised to GABA or acetylated to *N*-acetyl putrescine (Fig. 3C).(15–17) There was 300-fold less *N*-acetyl putrescine and 2.4-fold more GABA in SQ-grown *E. coli* (Fig. 3A), suggesting that more oxidation and less acetylation of putrescine occurs during sulfoglycolysis. The lower levels of *N*-acetylputrescine may indicate lower levels of acetyl-CoA production.

**Fig 3.**
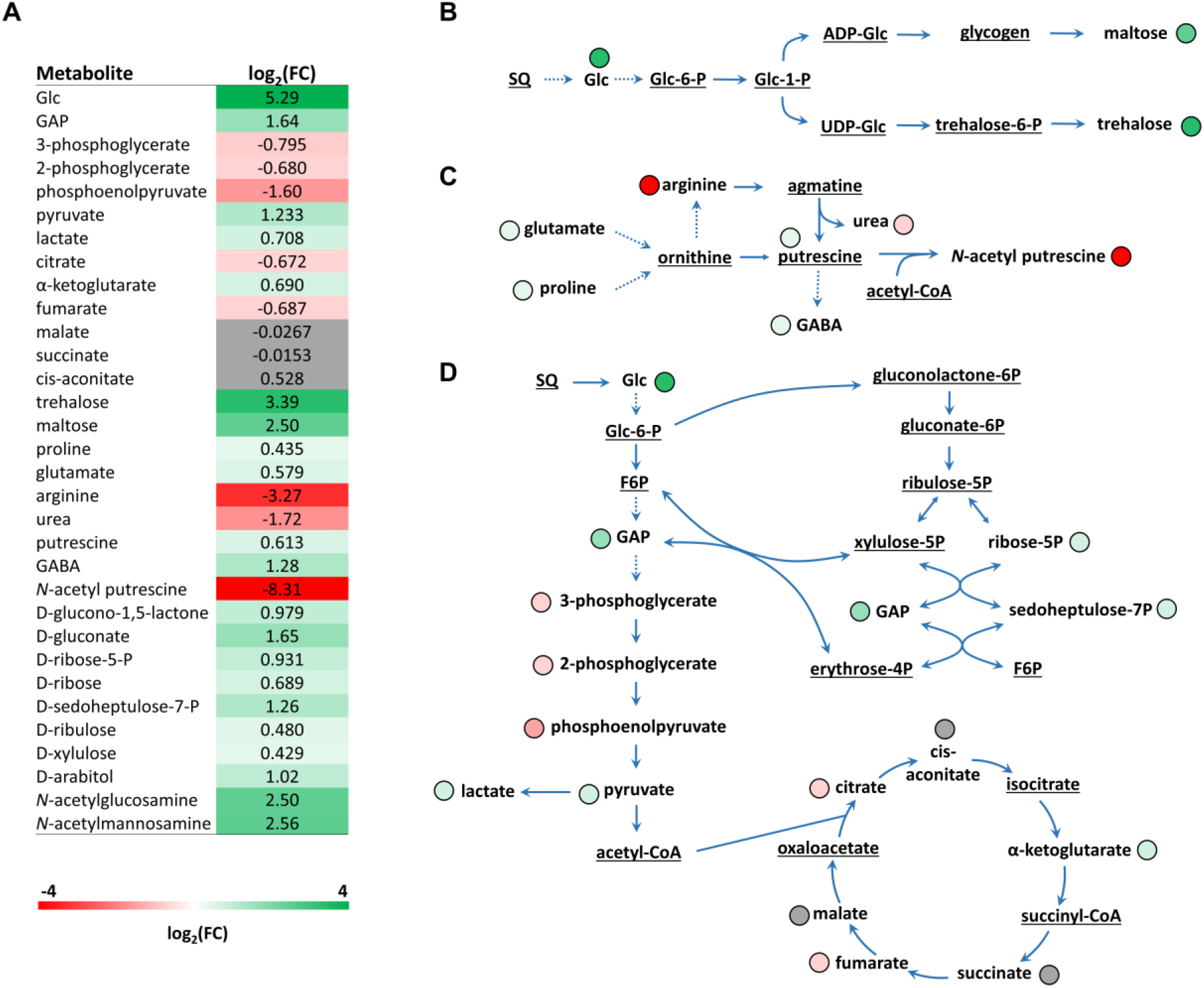
(A) Heat plot of selected metabolites detected in mid-log phase *E. coli* grown on Glc and on SQ depicting fold changes (FC). Selected metabolic pathways in *E. coli*(15–17) grown on minimal media, with FC indicated by coloured text: (B) trehalose, maltose and glycogen biosynthesis,(19, 20, 41, 42) (C) polyamine metabolism(18) and (D) central carbon metabolism. Metabolites highlighted in green are more abundant in SQ-grown *E. coli*, and those highlighted in red are less abundant. Metabolites shown in grey were not statistically different (FDR-adjusted p-value > 0.05). Metabolites shown in black were not detected.

### Sulfoglycolysis perturbs purine and pyrimidine metabolic pools

Adenosine was approximately 6-fold less abundant and xanthine >360-fold less abundant in SQ-grown versus Glc-grown *E. coli* (Table S1), indicative of a perturbation to purine metabolism. Pyrimidine metabolism was also perturbed: in SQ-grown *E. coli*, there was >95-fold less orotic acid, as well as less of the orotic acid precursors *N*-carbamoyl-L-aspartate (>100-fold less) and dihydroxyorotate (230-fold less) (Table S1). In *E. coli* BW25113, orotic acid is a metabolic end product;(22) thus, the lower levels of orotic acid may suggest decreased input into pyrimidine metabolism in sulfoglycolytic *E. coli*. Consistent with this observation, broader effects on pyrimidine metabolism were evident: there was 1.3-fold less uracil and 9-fold less of the uracil degradation product β-alanine in sulfoglycolytic *E. coli*, as well as 2.7-fold more thymine. The cytosine pool was not significantly perturbed (Table S1).

### Pools of redox mediators are perturbed upon switching to sulfoglycolysis

Glutathione and NAD(P)H help maintain the redox status of *E. coli* and are cofactors for a wide range of oxidoreductases. In SQ-grown *E. coli*, there was 7-fold more glutathione and 1.7-fold more nicotinate and nicotinamide versus glycolytic *E. coli* (Table S1), showing that sulfoglycolysis perturbs these key species in redox biochemistry.

### SQ-grown *E. coli* accumulate carbohydrate during mid-log phase

Various non-phosphorylated sugars (Glc, galactose, mannitol) and oligosaccharides (trehalose, maltose) accumulated in SQ-grown *E. coli* during mid-log phase (Fig. 2C), suggesting that a significant fraction of triose-P generated during SQ catabolism is diverted into synthesis of hexose phosphates (hexose-P) via the action of the reversible FBP aldolase and the dedicated gluconeogenic enzyme, fructose bisphosphatase (FBPase). The accumulation of Glc (>39-fold increase in SQ-fed versus Glc-fed bacteria) was unexpected and may reflect the activity of a hexose-P phosphatase (Fig. 2C, 3A) or constitutive cycling of glycogen pools via glycosidases as well as glycogen phosphorylase. The conversion of SQ to Glc via the gluconeogenic pathways was confirmed by labelling *E. coli* with (^13^C_6_)SQ (labelled:unlabelled 1:1). The presence of a prominent +3 isotopomer in the free Glc pool reflects the incorporation of triose-P into hexose synthesis.

Quantitative analysis reveals that there is more GAP but less phosphoenolpyruvate, 3-phosphoglycerate and 2-phosphoglycerate in mid-log phase SQ-grown *E. coli* (Fig. 3A). In contrast, higher levels of pyruvate and lactate were present in SQ-grown *E. coli*, which may indicate slower consumption and/or production of these metabolites through degradative pathways during adaptation to sulfoglycolysis. Similarly, the TCA cycle was also perturbed by the switch from growth on Glc to SQ. α-Ketoglutarate was 1.6-fold more abundant and fumarate and citrate 1.6-fold less abundant in the SQ-grown *E. coli* (Fig. 3A). There were no statistically significant differences in the malate, succinate and cis-aconitate pools. The perturbations to the TCA cycle and lower glycolytic metabolites are consistent with diversion of a significant portion of triose-P produced by sulfoglycolysis into gluconeogenesis, away from these downstream metabolic pathways.

The PPP metabolites D-glucono-1,5-lactone, D-gluconate, D-ribose-5-phosphate, D-ribose, D-seduheptulose-7-phosphate, D-ribulose, D-xylulose and D-arabitol were between 1.5-fold and 3-fold more abundant in SQ-grown *E. coli* (Fig. 3A), consistent with a general increased flux into sugar-phosphate synthesis under these growth conditions. *N*-Acetylglucosamine and *N*-acetylmannosamine were also >5-fold more abundant in SQ-grown *E. coli* (Fig. 3A). The accumulation of these sugars could reflect both increased flux into amino-sugar synthesis, as well as reduced rate of utilization of these sugars for cell wall biosynthesis in slower-growing SQ-fed bacteria.(8)

### Sulfoglycolysis perturbs cell wall biosynthesis in *E. coli*

The levels of *N*-acetylglucosamine, a key component of the peptidoglycan cell wall and outer membrane of *E. coli*, (23, 24) and *N*-acetylmannosamine, which can interconvert with the former,(15–17) were >5-fold larger in log-phase SQ-grown *E. coli* (Fig. 3A). The pools of fatty acids, key precursors required for the lipopolysaccharide outer membrane,(25) were also perturbed; all detected fatty acid pools, except linoleic acid, were 1.4-fold to 1.9-fold larger in SQ-grown *E. coli* (Table S1). This may reflect the lower growth rate of SQ-grown *E*. coli,(8) and hence slower consumption of these cell wall precursors. Alternatively, while cell pellets harvested from Glc-grown and SQ-grown *E. coli* were of similar size, the SQ-grown pellets were more difficult to resuspend in the extraction solvent, which may reflect differences in cell wall structure of glycolytic and sulfoglycolytic *E. coli*, such as thicker cell walls and/or cell walls that lead to greater cell adhesion.

### *E. coli* switch from sulfoglycolysis to glycolysis during stationary phase adaptation

The finding that mid-log phase *E. coli* accumulate trehalose and maltose, as well as intermediates in the PPP and amino-sugar synthesis when grown on SQ, indicated that a substantive proportion of triose-P is diverted into hexose/pentose phosphate synthesis via the final steps of gluconeogenesis or the PPP transketolase/transaldolase enzymes (Fig. 2C, 3A, Table 1). To assess whether growth on SQ also resulted in the accumulation of glycogen, the major *E. coli* carbohydrate reserve,(26–28) Glc- and SQ-grown *E. coli* were collected at five points along the growth curve and differentially extracted to recover low molecular weight oligosaccharides and glycogen. Maltose and trehalose were recovered in the methanol/water extract, and total protein and glycogen recovered in the insoluble pellet. The total carbohydrate content of the soluble fraction was also quantitated after methanolysis and conversion of Glc and fructose to their TMS derivatives. Storage carbohydrate content was normalized to total protein content for analysis (Fig. 4, S1, S2).

**Fig 4.**
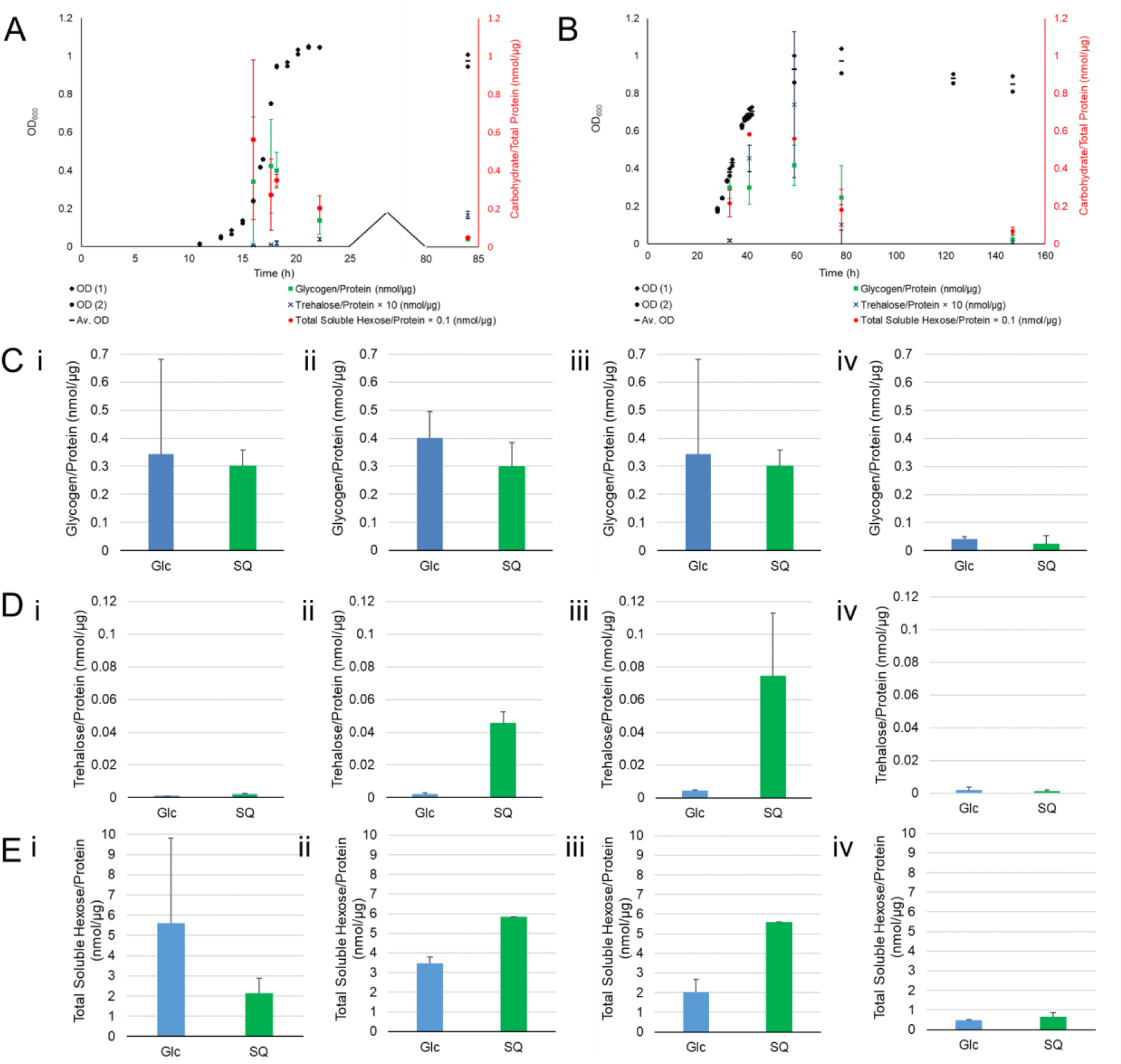
Storage carbohydrate content of Glc-grown and SQ-grown *E. coli* across the growth curve. (A) Storage carbohydrate content overlaid on growth curve of Glc-grown *E. coli*, (B) storage carbohydrate content overlaid on growth curve of SQ-grown *E. coli*, (C) glycogen content (measured as total Glc present in insoluble fraction) of Glc-grown and SQ-grown *E. coli*, (D) trehalose content of Glc-grown and SQ-grown *E. coli* and (E) Glc content of soluble fraction of Glc-grown and SQ-grown *E. coli*. For (C), (D) and (E): (i) early log phase, (ii) transition to stationary phase, (iii) early stationary phase and (iv) late stationary phase. Data shown is per cell pellet and is average of two independent replicates (mean ± SD, *n* = 2) with exception of data for SQ shown in (E) (ii) and (E) (iii) in which n = 1.

In Glc-grown *E. coli*, glycogen levels rose during log phase, peaking at the transition to stationary phase (or in late log phase), before declining as stationary phase progressed (Fig. 4A, 4C). This is broadly consistent with a previous report for this strain of *E. coli* (BW25113) in M9 minimal media containing 0.2% Glc (compared with 0.072% Glc in this work), in which glycogen levels peak around late log phase or the transition to stationary phase, and decline as stationary phase progresses.(29) SQ-grown *E. coli* also produced glycogen, accumulating during log phase, peaking in early stationary phase, then falling as stationary phase progressed (Fig. 4B, 4C). Peak levels were similar in SQ- and Glc-grown *E. coli*. Collectively, these data show that *E. coli* grown on SQ continue to accumulate glycogen and low molecular weight oligosaccharides at the expense of carbon flow into lower glycolysis. The accumulated glycogen appears to function as an alternative carbon source once extracellular Glc or SQ is depleted and cells enter stationary phase.

To verify that depletion of glycogen during stationary phase in SQ-grown *E. coli* was due to catabolism and not conversion to smaller storage carbohydrates such as the disaccharides trehalose/maltose, we analysed the carbohydrate content of the soluble fraction of stationary phase cells. No maltose was detected under either growth condition, and only very low levels of trehalose (an order of magnitude smaller than glycogen levels) were detected in SQ-grown *E. coli*, with levels decreasing as stationary phase progressed (Fig. 4B, 4C, 4D). Only trace amounts of trehalose were detected in Glc-grown *E. coli* during all stages of growth. In the total monosaccharide analysis, we targeted Glc and fructose. Only Glc was detected in both Glc- and SQ-grown *E. coli*, with levels an order of magnitude higher than glycogen in SQ-grown *E. coli* (Fig. 4B, 4C, 4E), indicating that most carbohydrate in Glc- and SQ-grown *E. coli* is present as free Glc and/or unidentified Glc-containing disaccharides. Glc levels rose across log phase in SQ-grown *E. coli*, then fell as stationary phase progressed. Taken together, these results suggest that glycogen accumulated during log-phase growth of *E. coli* on SQ is consumed upon transition to stationary phase, indicating a reversal in carbon flux from sulfoglycolytic/gluconeogenic to upper glycolytic.

## Discussion

Metabolism of SQ through the sulfo-EMP pathway leads to formation of triose-P, bypassing key steps in upper glycolysis that generate G6P and F6P for the PPP and the synthesis of cell wall intermediates. As a result, triose-P generated by SQ metabolism must be diverted into upper gluconeogenesis for hexose/pentose-phosphate production, as well as further catabolized in ATP-generating steps in lower glycolysis. This requirement imposes a double bioenergetic burden on SQ-grown bacteria: sulfoglycolysis only provides one 3C fragment and consumes one molecule of ATP. Furthermore, while upper glycolysis produces no reducing power, the sulfo-EMP pathway consumes one molecule of NADH (for reduction of SLA to DHPS) per SQ. While *E. coli* is thought to derive most of its ATP from oxidative phosphorylation when utilizing Glc as carbon source, the markedly reduced growth rate of SQ-grown *E. coli* compared to Glc-grown *E. coli*(8) indicates that the bioenergetic burden associated with SQ metabolism is substantial.

To partially compensate for the reduced carbon yield associated with SQ versus Glc metabolism, *E. coli* were cultivated in 8 mM SQ and 4 mM Glc. This ensured that the quantity of carbon (and potentially energy) available to the cells was essentially the same for both growth substrates. Nonetheless, substantially slower (approx. 5-fold) growth was observed for SQ-grown *E. coli* compared to Glc-grown *E. coli*. Comparative analysis of metabolite abundances in mid-log phase *E. coli* grown on SQ or Glc revealed that under SQ- or Glc-replete conditions there were large perturbations in levels of amino acids, various sugars and assorted cellular metabolites of a wide range of pathways, and possibly cell wall peptidoglycan. These changes are likely secondary effects that arise from large differences in the channelling of carbon that occurs from switching from growth on Glc (and use of both upper and lower glycolysis) to use of the sulfo-EMP pathway (use of sulfoglycolysis, lower glycolysis and upper gluconeogenesis); that is, these changes arise due to a switch from glycolytic to gluconeogenic metabolism.

Unexpectedly, we show that log phase sulfoglycolytic cells produce large quantities of hexoses and disaccharides, as well as the storage polysaccharide glycogen. Thus, under both Glc- and SQ-replete conditions, *E. coli* diverts some of the G6P produced through upper glycolysis/gluconeogenesis into storage carbohydrates. Glycogen is typically produced in *E. coli* when excess carbon is present but growth is limited by deficiency of an essential nutrient(s) required for growth.(26) Similarly, cultivation on SQ may lead to reduced flux into lower glycolysis and/or the TCA cycle and reduced synthesis of multiple anabolic intermediates needed for biomass (protein, nucleic acid, lipid) accumulation, leading to diversion of the resultant excess triose-P into carbohydrate synthesis.

Alternatively, the unbalanced diversion of triose-P into hexose-P synthesis during SQ metabolism may lead to reduced flux into lower glycolysis, oxidative phosphorylation and downstream metabolic pathways. Reduced downstream flux will result in less ATP, reducing power and carbon-based building blocks being available in sulfoglycolytic *E. coli*, which may be the main origin of the adjustments undertaken by *E. coli* to maintain balanced growth and thus the lower rate of growth on SQ. In accordance with this analysis, products of the TCA cycle were perturbed, suggesting an adaptation to reduced carbon input, and putrescine acetylation was depressed, potentially due to a lack of acetyl-CoA (Fig. 1C, 3C). Lower levels of pyrimidine biosynthesis metabolites may also arise from a deficiency of carbon building blocks. Amino acid levels were generally lower in sulfoglycolytic cells, which could reflect the overall energy state of the cell, given that protein synthesis is one of the most energy-intensive cellular process (Fig. 2B, 2C). Similarly, in SQ-grown *E. coli*, putrescine biosynthesis appears to prefer the more energetically favourable pathway direct from glutamate and/or proline via ornithine, with the alternative pathway from arginine via agmatine seemingly disfavoured (Fig. 3C).(15–18) Levels of glutathione and other redox-active metabolites were perturbed and oxidation of putrescine to GABA appeared to increase during sulfoglycolytic metabolism, potentially reflecting an altered redox state.

Organisms regulate levels of intracellular metabolites to prevent their accumulation to toxic levels. This can be especially pronounced during logarithmic growth, when metabolite production can far exceed cellular needs, leading to spillover. For example, yeast possess an energy-consuming futile cycle that consumes ATP through the production of trehalose from the sugar nucleotide UDP-Glc and Glc, while no ATP is produced in the hydrolysis of trehalose to Glc, even under conditions such as heat-shock where trehalose is accumulated.(30) Formation of glycogen in SQ-grown *E. coli* is a resource-intensive process: formation of one G6P from triose-P through gluconeogenesis requires investment of two ATP and consumption of two SQ; UTP must also be invested to form UDP-Glc. In Glc-grown *E. coli*, diversion of Glc into glycogen via G6P is less energy-intensive, requiring investment of only one ATP and one UTP. Futile cycling of cellular glycogen and other storage polysaccharides provides a mechanism to consume overflow ATP via conversion of G1P to UDP-Glc to make glucosidic linkages, and then release this carbon as G1P by phosphorolysis. Possibly, accumulation of cellular glycogen occurs to provide a substrate for an energy-consuming futile cycle.

Analysis of the storage polysaccharide content of *E. coli* cells grown on both Glc and SQ at different stages of growth indicates that upon transition to stationary phase, carbohydrates stored during log phase are consumed. In the case of SQ-grown *E. coli*, depletion of SQ appears to trigger a reverse diauxic shift whereby cells switch from a limited gluconeogenic metabolism to a canonical glycolytic metabolism as Glc/G6P is mobilized from glycogen/trehalose/maltose breakdown. Thus, growth on SQ requires multiple changes in the direction of flux of central carbon metabolism depending on growth state. The requirement for a change in flux from the glycolytic to gluconeogenic direction upon the switch to growth on SQ may in part contribute to the slow adaptation of *E. coli* to SQ;(8) in *E. coli*,switching from glycolytic to gluconeogenic metabolism also induces a long lag phase.(31, 32) In contrast, in *E. coli*, the switch from gluconeogenic to glycolytic metabolism occurs quickly(31, 32) and so the reversal upon SQ exhaustion is expected to be rapid.

Most enzymes in upper glycolysis/gluconeogenesis are reversible, and glycolysis/gluconeogenesis is almost thermodynamically neutral. As such, only limited changes in protein expression are required for change in flux direction, assisted by the presence of concentration gradients as one substrate is depleted and another becomes dominant, as well as allosteric effects. Growth on Glc provides a strong gradient promoting flux in the glycolytic direction, while growth on SQ leads to production of large amounts of triose-P, reversing the concentration gradient and providing a driving force for upper gluconeogenesis. Expression of class I FBP aldolase of *E. coli*, which is induced by growth on gluconeogenic substrates and presumed to be utilized for gluconeogenesis (as opposed to the constitutively expressed class II aldolase presumed to be utilized for glycolysis),(33) may be required upon a switch to SQ. In contrast, it has been shown that switching of *E. coli* from growth on Glc to the gluconeogenic substrate acetate results in just 2-fold change in expression levels of the upper gluconeogenic enzyme FBPase,(32) a key regulatory enzyme of gluconeogenesis.(34) FBPase is subject to complex allosteric regulation, with AMP and G6P serving as allosteric inhibitors(34, 35) while TCA cycle intermediates (such as citrate and isocitrate),(36) phosphorylated 3C carboxylic acids (such as PEP(37) and 3PG(36)), sulfate(37) and phosphate(36) are allosteric activators. The key allosteric interactions appear to be activation by citrate and PEP(34, 36) and synergistic inhibition by AMP and G6P.(34) There is likely an interplay of the production of storage carbohydrates from hexose-P generated by gluconeogenesis in SQ-grown *E. coli*, and the inhibitory effect of G6P on FBPase.

## Conclusions

We provide evidence that growth of *E. coli* on SQ imposes a significant bioenergetic cost on the bacteria, as a result of the lower yield of carbon from each molecule of SQ catabolized, and the need to divert a significant fraction of triose-P produced into hexose/pentose-phosphate synthesis. As a result, the reduced rate of growth of *E. coli* on SQ likely stems from reduced flux of carbon into the TCA cycle and downstream metabolism, producing lower levels of carbon building blocks, NADH and ATP, thereby triggering large-scale changes to cell metabolism to maintain balanced growth. Further work is needed to define the factors that regulate the partitioning of carbon between upper gluconeogenesis and lower glycolysis under SQ-grown conditions, while the mobilization of low and high molecular weight carbohydrates following depletion of SQ indicates that the accumulation of these molecules is an important adaptive response. This work highlights the ascetic nature of growth on SQ and provides insights into the metabolic adaptations of *E. coli* to growth on this widespread but poorly studied organosulfur sugar.

## Material and Methods

### Reagents

SQ and (^13^C_6_)SQ synthesized according to methods described previously.(38)

#### Comparative metabolite profiling of *E. coli* on Glc and SQ

*E. coli* BW25113 (adapted to growth on the relevant substrate) was used to inoculate a 5 mL starter culture of M9 minimal media containing 4 mM Glc or a 3 mL starter culture of M9 minimal media containing 8 mM SQ. The Glc starter culture was grown to OD_600_ 0.0680 (6 h 20 min) and the SQ starter culture was grown to OD_600_ 0.1864 (41 h) at 37°C with shaking (250 rpm) and used to inoculate the six experimental cultures. These experimental cultures contained either 4 mm Glc or 8 mM SQ. The culture volume was 50 mL for Glc and 20 mL for SQ and the inoculant volume was 298 μL for Glc and 118 μL for SQ. The Glc cultures were grown at 30°C with shaking (250 rpm) for 11 h, then at 37°C with shaking (250 rpm) until mid-logarithmic phase was achieved (approx. OD_600_ 0.50). The SQ cultures were grown at 37°C with shaking (250 rpm) until mid-logarithmic phase was achieved (approx. OD_600_ 0.44). One 5 mL aliquot was harvested per culture.

Aliquots of cell culture media were metabolically arrested by infusing with ice-cold PBS (3 × aliquot volume) and placing in an ice-water slurry for 10 min. The cell suspension was centrifuged (4000 rpm, 10 min, rt) and the supernatant discarded. Cells were washed 3 times by resuspending them in ice-cold PBS, pelleting them by centrifugation (14000 rpm, 1 min, rt) and discarding the supernatant. Cell pellets were then centrifuged again (14000 rpm, 1 min, rt) to remove residual culture medium and PBS prior to metabolite extraction.

#### Extraction, derivatization and analysis of metabolites for comparative metabolite profiling

Cell pellets were resuspended in chilled extraction solution (500 μL) comprised of 3:1 MeOH: H_2_O and (^13^C_5_, ^15^N)valine (1 mM, 0.5 μL) and (^13^C_6_)sorbitol (1 mM, 0.5 μL) as internal standards. The suspensions were subjected to freeze-thaw cycles to facilitate lysis of the cells (30 s in liquid N_2_, 30 s in a dry ice/EtOH bath for 10 cycles), and shaken (9000 rpm, 10 min, 2°C). The samples were then centrifuged (12700 rpm, 5 min, 1°C) to remove cell debris and precipitated macromolecules.(13)

The cell lysate supernatant was transferred into glass inserts and dried in a rotational vacuum concentrator. To remove all residual H_2_O, all samples were washed with MeOH (50 μL). The glass inserts were transferred to 2 mL autosampler vials. Online derivatization was conducted using an autosampler robot (Shimadzu AOC6000). All samples were methoximated with methoxylamine hydrochloride solution (30 mg/mL in pyridine, 20 μL) for 2 h at 37°C, followed by trimethylsilyation in BSTFA + 1% TMCS (20 μL) for 1 h at 37°C with continuous mixing. Samples were incubated at rt for 1 h prior to GC-MS analysis.

Metabolite profiles were acquired on a Shimadzu 2010 GC coupled to a TQ8040 QqQ mass spectrometer. The inlet temperature was held at 280°C and helium was used as a carrier gas (purge flow = 5.0 mL/min, column flow = 1.1 mL/min). 1 μL of derivatized sample was injected into the GC-QqQ-MS in splitless mode. Chromatographic separation was achieved using a J&W DB-5 capillary column(14) (30 m × 0.25 mm × 1.00 μM). The oven had a starting temperature of 100°C, which was held for 4 min, then ramped to 320°C at 10°C/min and held for 10 min. The transfer line temperature was 280°C and the ion source temperature was 200°C. Compounds were fragmented using electron (EI) ionization. Argon was used as the collision-induced dissociation gas. The metabolite detection was performed using the Shimadzu Smart MRM database, which contains up to 521 targets, representing ≈ 350 metabolites with two multiple reaction monitoring (MRM) transitions (quantifier and qualifier) per target, including precursor ion, product ion, collision energy, retention index and time, with a minimal dwell time of 2 msec set up for the acquisition method. The Automatic Adjustment of Retention Time (AART) in GCMSsolution software (V 4.42, Shimadzu) and a standard alkane series mixture (C7-C33 Restek) were used to correct retention time shifts in the acquisition method when the column is cut or replaced.

#### Data analysis for comparative metabolite profiling

Using Shimadzu Lab Solutions Insight, each target was visually inspected and manually integrated if required. Targets that were lacking in either the qualifier or quantifier MRM or had a quantifier < 50% of the abundance of the qualifier or had inconsistent retention times (either between the qualifier and quantifier, or between samples) were discarded. This provided a data matrix for downstream data analysis, which was conducted using the Metaboanalyst software (https://www.metaboanalyst.ca/MetaboAnalyst/home.xhtml). Data was normalized by median and log-transformed. FDR-adjusted p-values were considered significant < 0.05.

KEGG Mapper Search & Colour Pathway (https://www.genome.jp/kegg/tool/map_pathway2.html, reference organism: *E. coli* K12 W3110) was used to assist in drawing conclusions. In cases where more than one target corresponding to a single metabolite was detected, the fold changes of the targets were averaged to give the fold change for the metabolite.

#### Metabolite analysis for *E. coli* grown on (^13^C_6_)Glc and (^13^C_6_)SQ

*E. coli* BW25113 (adapted to growth on the relevant substrate) was used to inoculate a 5 mL starter culture of M9 minimal media containing 4 mM Glc or a 3 mL starter culture of M9 minimal media containing 8 mM SQ. The Glc starter culture was grown to OD_600_ 0.0808 (8 h) and the SQ starter culture was grown to OD_600_ 0.1964 (41 h) at 37°C with shaking (250 rpm) and used to inoculate the experimental cultures. These experimental cultures were either unlabelled (4 mM (^12^C_6_)Glc or 8 mM (^12^C_6_)SQ), labelled for the entire growth period (2 mM (^12^C_6_)Glc + 2 mM (^13^C_6_)Glc or 4 mM (^12^C_6_)SQ + 4 mM (^13^C_6_)SQ) or labelled for a portion of the growth period (4 mM (^12^C_6_)Glc, resuspended into 2 mM (^12^C_6_)Glc + 2 mM (^13^C_6_)Glc 2.75 h before harvest; or 8 mM (^12^C_6_)SQ, resuspended into 4 mM (^12^C_6_)SQ + 4 mM (^13^C_6_)SQ 4 h before harvest). The culture volume was 50 mL for Glc and 20 mL for SQ and the inoculant volume was 200 μL for Glc and 112 μL for SQ. The Glc cultures were grown at 30°C with shaking (250 rpm) for 11 h, then at 37°C with shaking (250 rpm) until mid-logarithmic phase was achieved (approx. OD_600_ 0.50). The SQ cultures were grown at 37°C with shaking (250 rpm) until mid-logarithmic phase was achieved (approx. OD_600_ 0.44). Three 5 mL aliquots were harvested per culture.

Cell pellets were resuspended in chilled extraction solution (500 μL) comprised of 3:1 MeOH: H_2_O and *scyllo*-inositol (1 mM, 0.5 μL) as an internal standard. Cell suspensions were subjected to freeze-thaw cycles to facilitate lysis of the cells (30 s in liquid N_2_, 30 s in a dry ice/EtOH bath for 10 cycles), and shaken (9000 rpm, 10 min, 4°C). The samples were then centrifuged (14000 rpm, 5 min, 1°C) to remove cell debris.(13)

The cell lysate supernatant was transferred into glass inserts and dried in a rotational vacuum concentrator. To remove all residual H_2_O, all samples were washed with MeOH (50 μL). The glass inserts were transferred to 2 mL autosampler vials. Online derivatization was conducted with a Gerstel MSP2 XL autosampler robot (Gerstel, Germany). All samples were methoximated with methoxylamine hydrochloride solution (30 mg/mL in pyridine, 20 μL) for 2 h at 37°C, followed by trimethylsilyation in BSTFA + 1% TMCS (20 μL) for 1 h at 37°C with continuous mixing. Samples were incubated at rt for 1 h prior to GC-MS analysis.

Metabolite profiles were acquired on an Agilent 7890A Gas Chromatograph coupled to a 5975 mass spectrometer as a detector. 1 μL of derivatized sample was injected in splitless mode into a split/splitless inlet set at 250°C. Chromatographic separation was achieved using a J&W VF-5 ms capillary column (30 m × 0.25 mm × 0.25 μM + 10 m duraguard). The oven had a starting temperature of 35°C, which was held for 1 min, then ramped to 320°C at 25°C/min and held for 5 min. Helium was used as the carrier gas at a flow rate of 1 mL/min. Compounds were fragmented using electron impact (EI) ionization and detected across a mass range of 50-600 amu with a scan speed of 9.2 scans/s.(39)

Chromatograms produced from unlabelled samples were used for pool size comparisons. Using Agilent’s Mass Hunter Quantitative Analysis software for GC-MS, metabolites contained within an in-house Metabolomics Australia library, with the addition of some metabolites detected and identified in the chromatograms using Agilent ChemStation for GC-MS and the Fiehn L metabolite library, were identified and representative target ion areas were integrated. Each detected metabolite in each chromatogram was visually inspected and manually integrated if required. These formed an output data matrix for downstream data analysis,(39) which was conducted using the Metaboanalyst software (http://www.metaboanalyst.ca/faces/ModuleView.xhtml). Data was normalized by median and log-transformed. P-values were considered significant < 0.05 and fold changes were considered significant < 1.5.

To detect labelled metabolites, chromatograms were processed using the Non-targeted Tracer Fate Detection software (https://ntfd.bioinfo.nat.tu-bs.de/) (peak threshold: 5, minimum peak height: 5, deconvolution width: 5 scans, minimum % label: 5%, maximum % label: 100%, minimum R^2^: 0.95, maximum fragment deviation: 0.20, required no. of labelled fragments: 1, M1 correction: 0.0109340), followed by manual verification by visual inspection using Agilent ChemStation for GC-MS. Identification was achieved by comparison with the unlabelled samples, from which metabolites were identified using Agilent ChemStation for GC-MS and the Fiehn L metabolite library.

#### Storage carbohydrate analysis

*E. coli* BW25113 (adapted to growth on the relevant substrate) was used to inoculate a 5 mL starter culture of M9 minimal media containing 4 mM Glc or a 3 mL starter culture of M9 minimal media containing 8 mM SQ. The Glc starter culture was grown to OD_600_ 0.0630 (5 h 15 min) and the SQ starter culture was grown to OD_600_ 0.2524 (41 h) at 37°C with shaking (250 rpm) and used to inoculate the two experimental cultures that contained either 4 mm Glc or 8 mM SQ. The culture volume was 50 mL for Glc and 20 mL for SQ and the inoculant volume was 322 μL for Glc and 87 μL for SQ. The Glc cultures were grown at 30°C with shaking (250 rpm) for 11 h, then at 37°C with shaking (250 rpm). The SQ cultures were grown at 37°C with shaking (250 rpm). Bacteria were harvested at 5 different time points, with two aliquots harvested per culture at each time point. The harvest points for the Glc cultures were: 960 min (OD_600_ 0.2380, 0.24120), 1060 min (OD_600_ 0.7528, 0.7504), 1090 min (OD_600_ 0.9512, 0.9436), 1330 min (OD_600_ 1.0444, 1.0472), and 5040 min (OD_600_ 1.0100, 0.9464). The harvest points for the SQ cultures were: 33 h (OD_600_ 0.3988, 0.3636), 41 h (OD_600_ 0.7172, 0.6736), 59 h (OD_600_ 1.0012, 0.8600), 78 h (OD_600_ 1.0376, 0.9088) and 147 h (OD_600_ 0.8920, 0.8100). The culture volume harvested contained the same number of cells (by OD_600_) as 500 μL of OD_600_ 0.5 culture.

Aliquots of cell culture media were metabolically arrested by infusing with ice-cold PBS (800 μL, except for 960 min harvest for Glc, and 33 h harvest for SQ, which used 400 μL and 700 μL respectively) and placing in an ice-water slurry for 5 min. The cell suspension was centrifuged (14000 rpm, 5 min, rt) and the supernatant discarded. Cells were washed 3 times by resuspending them in ice-cold PBS (0.2 mL), pelleting them by centrifugation (14000 rpm, 1 min, rt) and discarding the supernatant. Cell pellets were then centrifuged again (14000 rpm, 1 min, rt) to remove residual culture medium and PBS prior to glycogen extraction. Cell pellets were thawed in 100 μL of MilliQ water. 375 μL of 1:2 CHCl3: MeOH was then added. The samples were incubated at rt for 1 h and vortexed regularly. The samples were then centrifuged (15000 rpm, 10 min, rt) to separate soluble and insoluble metabolites.

#### Analysis for glycogen content

Insoluble metabolites were resuspended in 100 μL of MilliQ water. 10 μL of each sample was transferred to a glass insert. Each insert also contained 1 nmol of *scyllo*-inositol. Standards were also prepared: 1 nmol *scyllo*-inositol only, 5 nmol Glc, 2.5 nmol Glc, 1 nmol Glc, 0.5 nmol Glc and 0.25 nmol Glc. All inserts were dried in a rotational vacuum concentrator (1 h). 50 μL of 2 M TFA in water was then added, and the samples and standards incubated for 2 h at 100°C and dried under N_2_ (1 h). 10 μL of MeOH was added and the inserts dried in a rotational vacuum concentrator to remove all H_2_O (25 min). 50 μL of 0.5 M HCl in MeOH was added and the samples and standards incubated for 17 h at 80°C. 10 μL pyridine was added to neutralize the HCl and the inserts were dried in a rotational vacuum concentrator (30 min). 20 μL of pyridine was added, followed by 20 μL of BSTFA + 1% TMCS.

Samples and standards were analysed on an Agilent 7890A Gas Chromatograph coupled to a 5975 mass spectrometer as a detector. The inlet temperature was 250°C and helium was used as a carrier gas (purge flow = 20 mL/min, column flow = 1 mL/min). 1 μL of derivatized sample was injected. Chromatographic separation was achieved using a DB-5 capillary column (30 m × 0.25 mm × 1.00 μM). The oven had a starting temperature of 70°C, which was held for 1 min, then ramped to 295°C at 12.5°C/min, then 320°C at 25°C/min. The transfer line temperature was 250°C and the ion source temperature was 230°C. Compounds were fragmented using electron (EI) ionization. Spectra were acquired over the range 50-500 m/z.

Data was analysed using the Agilent MassHunter software. The total ion chromatogram for each sample and standard was integrated. Sugars were identified based on GC retention time and mass spectra of authentic standards. The ratio of the minor β-anomer of Glc (16.4 min) to *scyllo*-inositol (18.7 min) was determined, with that peak chosen as it was well resolved compared to the major α-anomer. A Glc calibration curve was constructed using the β-Glc: *scyllo*-inositol ratio from the 5 Glc standards and used to determine the quantity of Glc in each of the samples. The quantity of Glc found in the *scyllo*-inositol blank was subtracted, and this value used to determine the quantity of Glc in each cell pellet.

#### Analysis for disaccharide content(40)

120 μL of the soluble fraction of each sample and 1 nmol *scyllo*-inositol was transferred to a glass insert. Standards were prepared for trehalose and maltose at the following quantities: 100 nmol, 50 nmol, 25 nmol, 12.5 nmol, 6.25 nmol, 3.125 nmol, 1.563 nmol, 0.781 nmol, 0.391 nmol and 0.195 nmol. All inserts were dried in a rotational vacuum concentrator (1 h). 20 μL of 20 mg/mL methoxyamine hydrochloride in pyridine was added and the samples and standards incubated with continuous mixing for 14 h at 25°C. 20 μL of BSTFA + 1% TMCS was added and samples and standards incubated for 1 h at 25°C.

Samples and standards were analysed on an Agilent 7890A Gas Chromatograph coupled to a 5975 mass spectrometer as a detector. The inlet temperature was kept at 250°C. Chromatographic separation was achieved using an Agilent CP9013 VF-5ms column (30 m × 0.25 mm × 0.25 μM) with 10 m EZ-guard. The oven had a starting temperature of 70°C, which was held for 2 min, then ramped to 295°C at 12.5°C/min, then 320°C at 25°C/min, then held at 320°C for 3 min. The transfer line temperature was 280°C. Compounds were fragmented using electron (EI) ionization.

Data was analysed using the Agilent MassHunter software. The total ion chromatogram for each sample and standard was integrated. Sugars were identified based on GC retention time and mass spectra of authentic standards. The ratio of trehalose (19.8 min) to *scyllo*-inositol (15.1 min) was determined. Calibration curves were constructed using the disaccharide: *scyllo*-inositol ratio from 7 of the standards (25 nmol, 50 nmol and 100 nmol excluded due to loss of linearity) and used to determine the quantity of trehalose in each of the samples. This value was used to determine the quantity of trehalose in each cell pellet.

#### Monosaccharide derivatization and analysis(40)

80 μL of the soluble fraction of each sample was transferred to a glass insert that also contained 1 nmol *scyllo*-inositol. Standards were also prepared for glucose and fructose at the following quantities: 100 nmol, 50 nmol, 25 nmol, 12.5 nmol, 6.25 nmol, 3.125 nmol, 1.563 nmol, 0.781 nmol, 0.391 nmol and 0.195 nmol. All inserts were dried in a rotational vacuum concentrator (1 h). 50 μL of 0.5 M HCl in MeOH was added and the samples and standards incubated for 3 h at 80°C. 10 μL pyridine was added to neutralize the HCl and the inserts were dried in a rotational vacuum concentrator (30 min). 20 μL of BSTFA + 1% TMCS was added and samples and standards incubated for 1 h at 25°C.

Samples and standards were analysed on an Agilent 7890A Gas Chromatograph coupled to a 5975 mass spectrometer as a detector. The inlet temperature was kept at 250°C. Chromatographic separation was achieved using an Agilent CP9013 VF-5ms column (30 m × 0.25 mm × 0.25 μM) with 10 m EZ-guard. The oven had a starting temperature of 70°C, which was held for 2 min, then ramped to 295°C at 12.5°C/min, then 320°C at 25°C/min, then held at 320°C for 3 min. The transfer line temperature was 280°C. Compounds were fragmented using electron (EI) ionization.

Data was analysed using the Agilent MassHunter software. The total ion chromatogram for each sample and standard was integrated. Sugars were identified based on GC retention time and mass spectra of authentic standards. Due to high variation in internal standard (*scyllo*-inositol) peak areas across the standards and samples, peak areas were normalized to the lowest detected internal standard peak area. The ratio of the normalized peak area of the first-eluting Glc peak (13.96 min) to the area of the *scyllo-*inositol peak (15.1 min) was determined. Calibration curves were constructed using the Glc: *scyllo*-inositol ratio from 4 of the standards (6.25 nmol – 50 nmol) and used to determine the quantity of Glc in each of the samples. This value used to determine the quantity of Glc in each cell pellet.

#### BCA Protein Assay

60 μL of insoluble metabolites suspended in MilliQ water from each sample was dried in a rotational vacuum concentrator (1.5 h, 50°C). 60 μL of 0.5% SDS in dH_2_O was added, the samples were vortexed, then boiled for 5 min and centrifuged (16100 rpm, 5 min, rt) to pellet cell debris. The supernatant and standards containing bovine serum albumin (BSA) were analyzed using a BCA protein assay kit (Sigma).

## Acknowledgements

We thank Elizabeth King for helpful technical support.

## Author Contributions

SJW and MJM conceptualized research; JM, DD, EK and ES conducted research; JM, MJM and SJW analyzed the data; and JM, MJM and SJW wrote the paper.

